# HMMR/RHAMM recruits SACK1D/FAM83D-CK1α complex at the mitotic spindle to control spindle alignment

**DOI:** 10.1101/2025.06.23.661043

**Authors:** Tyrell N. Cartwright, Naveen K. Nakarakanti, Karen Dunbar, Luke J. Fulcher, Selina Bader, Nicola T. Wood, Thomas J. Macartney, Gopal P. Sapkota

## Abstract

The SACK1D/FAM83D-CK1α complex assembles at the mitotic spindle to orchestrate proper spindle positioning and error-free progression through mitosis. The full molecular picture of how this complex assembles and disassembles over the cell division cycle remains to be fully defined. Here, we show that HMMR is critical for SACK1D-CK1α complex formation at the spindle, co-localizes with the SACK1D-CK1α complex throughout mitosis, and is necessary for correct mitotic spindle alignment. We find that HMMR binds to the C-terminal α-helix of SACK1D, and this helix is also important for the mitotic interaction between SACK1D and CK1α. We demonstrate that HMMR binding stabilizes SACK1D. We map the mitotic hyperphosphorylation sites on SACK1D and show that this hyperphosphorylation signals the destruction of SACK1D upon mitotic exit. The destruction also requires the C-terminal α-helix of SACK1D, suggesting that hyperphosphorylation of SACK1D in mitosis potentially exposes the C-terminal degron sequences resident on SACK1D. This study provides key molecular insights into the assembly and fate of the HMMR-SACK1D-CK1α complex at the mitotic spindle.

## Introduction

Scaffold Anchor of CK1 (SACK1) domain-containing protein D (SACK1D; known previously as FAM83D or CHICA) is a microtubule-localized scaffold protein conserved in vertebrates that orchestrates proper spindle alignment, ensuring timely cell division (Fulcher *et al*, 2019). In mitosis, SACK1D directly associates with and recruits the Ser/Thr protein kinase CK1α to the mitotic spindle via its conserved SACK1 domain (previously known as domain of unknown function 1669, DUF1669) (Fulcher *et al*, 2018; Fulcher *et al*., 2019). We previously showed that CK1α kinase activity at the mitotic spindle is essential for correct spindle positioning and timely, error-free progression through mitosis, and that CK1α recruitment to the mitotic spindle is dependent on SACK1D (Fulcher *et al*., 2019). We and others have also shown that SACK1D is recruited to the mitotic spindle through its association with the microtubule-associated protein hyaluronan-mediated motility receptor (HMMR, aka RHAMM or CD168) (Connell *et al*, 2017; Dunsch *et al*, 2012; Fulcher *et al*., 2019). However, the precise molecular mechanisms by which HMMR binds to and recruits SACK1D to the mitotic spindle remain to be defined.

HMMR, first identified as a receptor for hyaluronic acid, has been implicated in a wide range of cancers with high expression correlated to poor outcomes. Several studies on hepatocellular carcinoma (Wu *et al*, 2024; Zhang *et al*, 2025), breast cancer (Shabir *et al*, 2024), lung adenocarcinoma (Yu *et al*, 2025), acute lymphoblastic leukaemia (Ramirez-Chiquito *et al*, 2025), and head and neck squamous cell carcinoma (Zhang *et al*, 2024) point to HMMR playing an important role in cancer cell proliferation. While some studies have linked this to hyaluronan signalling pathways, most studies have focussed on HMMR as a prognostic indicator and have not examined the underpinning molecular pathways that HMMR controls. Studies examining the molecular roles of HMMR have shown that it acts intracellularly in concert with other microtubule and centrosomal proteins to govern structural cytoskeletal, mitotic and meiotic processes, which directly affect cell proliferation, correct cell division cycle progression and microtubule organization (He *et al*, 2020). Indeed, HMMR was shown to regulate microtubule nucleation and spindle assembly in *Xenopus* (Groen *et al*, 2004). HMMR has been shown to play an important role in the polo-like kinase (PLK)-dependent spindle positioning pathway, and has been shown to restrict the activity of the kinesin Eg5 by modulating TPX2 recruitment (Chen *et al*, 2018; Connell *et al*., 2017). HMMR also plays a role outside of mitosis in regulating TPX2 localization in neurons (Chen *et al*, 2024). Some studies have highlighted the importance of HMMR in other aspects of proliferation and mitosis (Jiang *et al*, 2021; Zhang *et al*, 2023). Structurally, HMMR contains a leucine zipper motif at its C-terminus, which was shown to be important for centrosomal localization (Maxwell *et al*, 2003). The N-terminus of HMMR, on the other hand, is known to be important for microtubule binding, as deletion of the first 189 residues of the protein abolished its localisation to microtubules (Dunsch *et al*., 2012). Though these studies paint a clear picture of HMMR as an intracellular, cytoskeletal and mitotic factor, the regulation bestowed by HMMR to the molecular mechanism underpinning CK1α recruitment to the mitotic spindle remains undefined.

SACK1D and HMMR are both cell cycle regulated proteins. Both proteins are upregulated in G2/M and are degraded upon mitotic exit, temporally coincident with other critical mitotic proteins, although precisely how this is achieved is not known (Fulcher *et al*., 2019). Through the interaction with its substrate binder Cdc20, the anaphase-promoting complex/cyclosome (APC/C) E3 ligase complex is responsible for the initiation of anaphase onset through the destruction of key substrates including both Cyclin B1, the activation cofactor of cyclin-dependent kinase 1 (CDK1), which is responsible for a myriad of critical mitotic phosphorylation events, and securin, an inhibitor of separase, which is responsible for cohesion cleavage allowing sister chromatids to separate (McAinsh & Kops, 2023). Recruitment to the APC/C requires the recognition of one of several destruction sequence motifs on target proteins, such as the D-box (RxxLxxxxN), KEN and ABBA motifs (McAinsh & Kops, 2023). Given the precise temporal control of both SACK1D and HMMR expression, it is possible that these too are targets of the APC/C, and as such contain mitotic degrons known to facilitate their destruction at anaphase onset. Additionally, SACK1D undergoes hyperphosphorylation in mitosis leading to an apparent molecular weight shift by SDS-PAGE of ∼25 KDa and this requires CK1α kinase activity (Fulcher *et al*., 2019). However, the precise sites of mitotic hyperphosphorylation on SACK1D and what effect hyperphosphorylation has on SACK1D function or fate remain undefined. In this study, we investigate the molecular details underpinning precisely how HMMR recruits SACK1D-CK1α complex to the mitotic spindle. Finally, we examine mitotic hyperphosphorylation of SACK1D and its role in the post-mitotic degradation of SACK1D and HMMR.

## Results

### HMMR co-localizes with SACK1D-CK1α complex at the spindle throughout mitosis and is required for SACK1D-CK1α complex formation and correct mitotic spindle alignment

We and others have previously reported that SACK1D localizes to the mitotic spindle (Dunsch *et al*., 2012; Fulcher *et al*., 2019) and is required for the recruitment of CK1α to the mitotic spindle to ensure correct spindle alignment and timely mitosis (Fulcher *et al*., 2019). We monitored the subcellular distribution of SACK1D and CK1α in unperturbed asynchronous SACK1D*^GFP/GFP^* and CK1α*^mCherry/mCherry^* double knock-in U2OS cells (Fulcher *et al*., 2019), generated through CRISPR/Cas9 genome editing, with confocal immunofluorescence microscopy through various stages of the cell cycle (**Figure 1A**). Both SACK1D and CK1α display diffused expression in interphase cells. At the G2/M transition both proteins begin to coalesce around centrosomes. By late prophase/early prometaphase, when the centrosomes have separated, both proteins are strongly associated with newly forming spindle fibres protruding from the centrosomes. This association remains strong throughout spindle assembly and persists all the way until anaphase, where both proteins appear to dissociate from the spindle as chromosomes segregate (**Figure 1A**). Similarly, HMMR displays identical spatiotemporal localization to SACK1D and CK1α, except in non-mitotic cells, where HMMR also stains interphase microtubules (**Figure 1B**). Absence of immunofluorescence signal for HMMR in HMMR*^-/-^* U2OS cells, which were generated by CRISPR/Cas9 genome editing and confirmed by genomic sequencing (**EV1**), confirmed the specificity of the antibody used for immunofluorescence (**Figure 1B**). Furthermore, when we analysed mitotic SACK1D-CK1α complex formation by immunoprecipitating CK1α, we confirmed the absolute requirement for the presence of HMMR, as the CK1α IPs failed to pull down SACK1D in HMMR*^-/-^* cells (**Figure 1C**). Similarly, HMMR did not co-precipitate with CK1α in SACK1D*^-/-^*cells (**Figure 1C**). In WT cells, both SACK1D and HMMR co-precipitated with CK1α specifically in mitosis (**Figure 1C**). We further confirmed the requirement of HMMR in mitotic SACK1D-CK1α complex formation, as siRNA-mediated depletion of HMMR in SACK1D*^GFP/GFP^* and CK1α*^mCherry/mCherry^* double knock in U2OS cells completely abolished the mitotic spindle accumulation of the SACK1D-CK1α complex seen in control conditions (**Figure 1D**). Similarly, HMMR*^-/-^* U2OS cells failed to recruit SACK1D and CK1α to the mitotic spindle (**Figure 2A**). However, retroviral-mediated restoration of the two reported transcriptional variants of HMMR, namely the full-length HMMR-GFP and a shorter variant lacking exon 4 (HMMR ΔE4-GFP), both restored CK1α and SACK1D at the mitotic spindle (**Figure 2A**). Consistent with these observations, CK1α co-precipitated SACK1D and HMMR in mitotic HMMR*^-/-^* U2OS cells restored with either HMMR-GFP or HMMR ΔE4-GFP, but not in control HMMR*^-/-^* U2OS cells (**Figure 2B**). Finally, we wanted to determine the effect of HMMR loss on the spindle alignment phenotype, which we previously reported was disrupted upon the loss of SACK1D or CK1α from the spindle (Fulcher *et al*., 2019). We performed mitotic spindle alignment assays using confocal microscopy and found that WT U2OS cells on average have spindles parallel to the culture surface, whereas HMMR*^-/-^* U2OS cells have randomly rotated spindles during late prometaphase, suggesting a significant misalignment of mitotic spindles in HMMR^-/-^ cells (**Figure 2C**). This spindle misalignment phenotype observed in HMMR^-/-^ U2OS cells was corrected by restoration of HMMR-GFP or its ΔE4 variant (**Figure 2C**).

**Figure 1.**
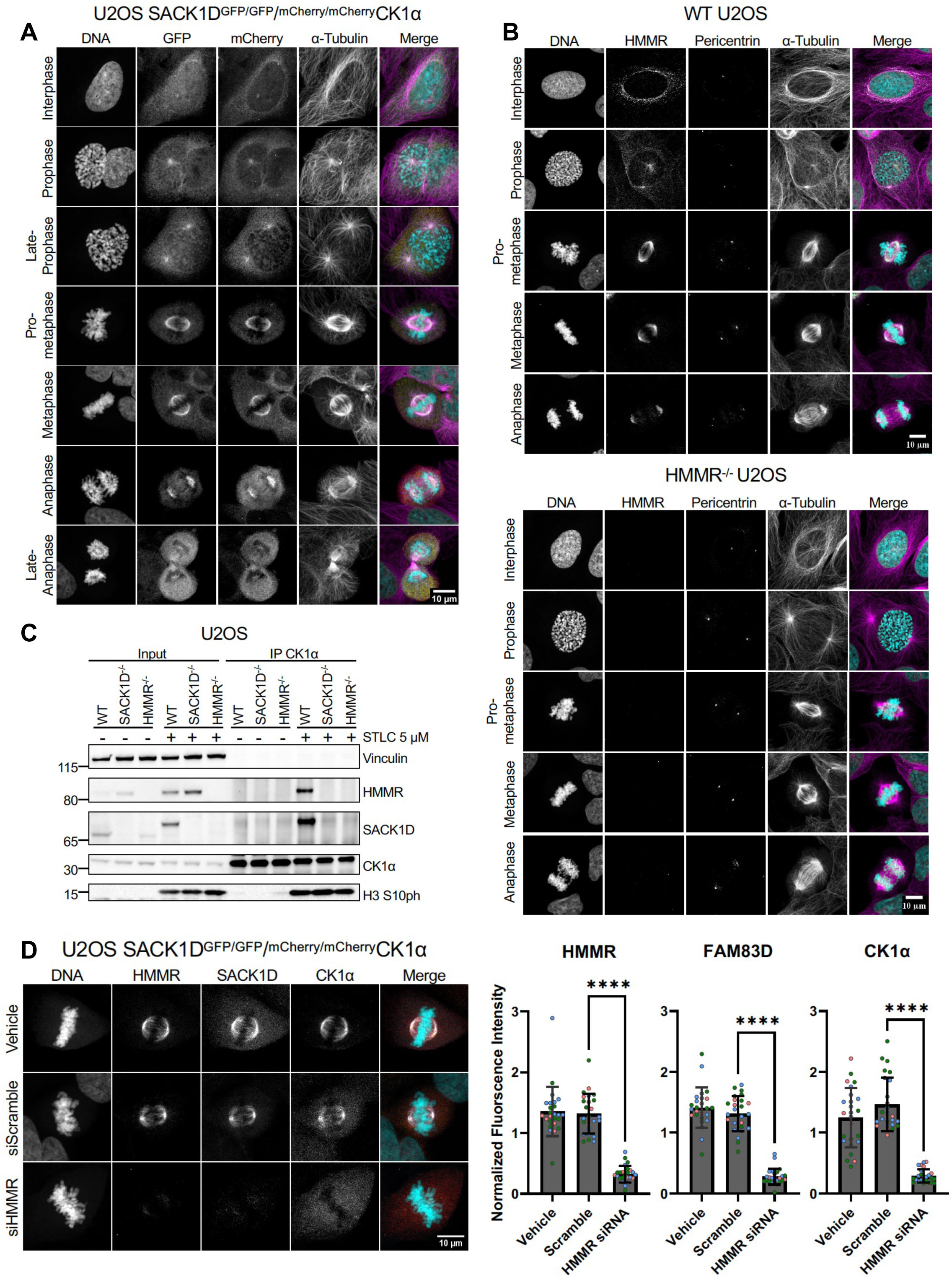
HMMR is a critical component of the SACK1D:CK1α spindle complex. **A-B**. Representative confocal immunofluorescence (IF) microscopy images of the SACK1D^GFP/GFP^/^mCherry/mCherry^CK1α double knock-in U2OS cells showing that SACK1D-GFP, mCherry-CK1α and HMMR all localize to the centrosomes and mitotic spindle throughout various stages of mitosis. **C**. Representative western blot with the indicated antibodies of lysate inputs and CK1α immunoprecipitates (IPs) in WT, SACK1D^-/-^ and HMMR^-/-^ U2OS cells under asynchronous conditions or those treated with STLC at 5 μM for 16 h to induce mitotic arrest. **D**. Representative confocal IF microscopy images of SACK1D^GFP/GFP^ and ^mCherry/mCherry^CK1α CRISPR double knock in U2OS cells treated with HMMR siRNA and stained with anti HMMR antibody. Graph indicates quantification of fluorescence intensity for each protein labelled in 3 independent experiments (n=25). Dots indicate individual cells with colours representing independent experiments. (**** p<0.001)

**Figure 2.**
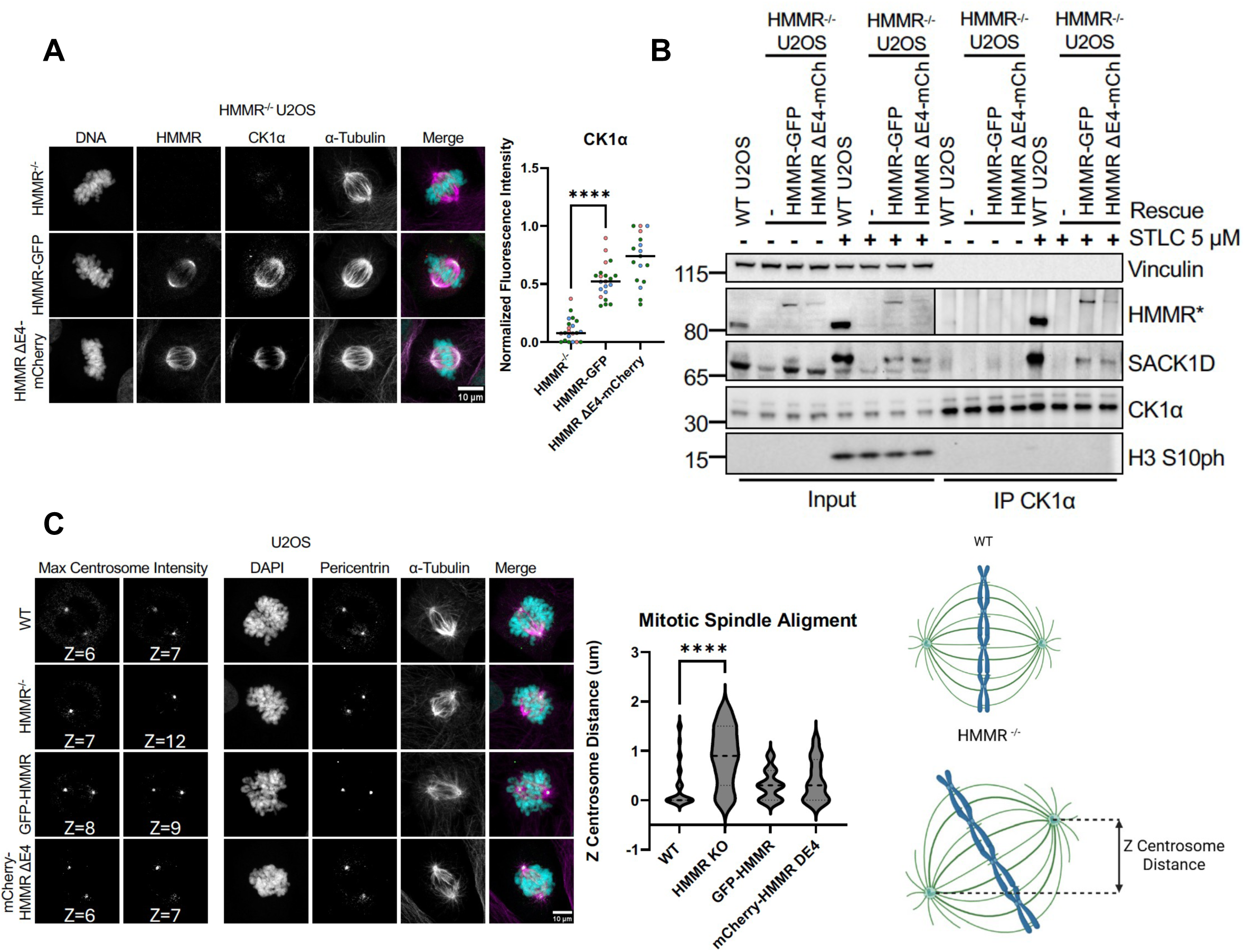
Restoration of HMMR in HMMR^-/-^ U2OS cells is sufficient to re-localize the SACK1D-CK1α complex to mitotic spindles and correct spindle alignment defects. **A**. Representative confocal IF microscopy images of HMMR^-/-^ U2OS cells either untransduced or transduced with retroviruses to stably express HMMR-GFP or HMMR-GFP lacking exon 4 (ΔE4-GFP) and stained for HMMR, CK1α and α-tubulin as indicated. Graph indicates quantification of fluorescence intensity for 3 independent experiments (n>=17). Dots indicate individual cells with colours representing each independent biological replicate (**** p<0.001). **B**. Representative western blot analysis with the indicated antibodies of lysate inputs and CK1α IPs in WT U2OS cells and HMMR^-/-^ U2OS cells stably expressing HMMR-GFP or HMMR-GFP lacking exon 4 (ΔE4-GFP). **C.** Confocal IF spindle alignment assay of WT U2OS cells, and HMMR^-/-^ U2OS cells that were either untransduced or transduced with retroviruses to stably express HMMR-GFP or ΔE4-GFP as indicated and stained for Pericentrin as a marker for centrosomes. Violin plot indicates data from three independent experiments (n>=24, **** p<0.001).

### The C-terminal α-helix of SACK1D is responsible for its association with HMMR as well as regulating its mitotic-specific association with CK1

To better understand how HMMR controls the role of the SACK1D-CK1α complex in ensuring proper spindle alignment, we set out to determine how SACK1D interacts with HMMR. Using AlphaFold3 (Abramson *et al*, 2024), the interaction between HMMR and SACK1D was predicted to involve the C-terminal α-helix of SACK1D and residues within the central coiled-coil region of HMMR (**EV2A&B**). Guided by the predicted structure, a series of mutations within the region of SACK1D predicted to disrupt HMMR binding or a truncated fragment (aa1-514) lacking the entire C-terminal α-helix were generated and their ability to interact with HMMR investigated (**Figure 3A**). SACK1D-GFP (WT-GFP), SACK1D*^M545A,L546A,M548A,L549A^*-GFP (2xML-GFP) and SACK1D*^1-514^*-GFP (1-514-GFP) were expressed in SACK1D*^-/-^* U2OS cells and immunoprecipitated using an anti-GFP nanobody immobilized on Sepharose beads. WT SACK1D-GFP was able to co-precipitate HMMR in both interphase and mitotic cells. However, neither 2xML-GFP nor 1-514-GFP were able to co-precipitate HMMR (**Figure 3B**), suggesting that the C-terminal α-helix of SACK1D is essential for mediating the interaction with HMMR. Another noteworthy observation is that the 2xML-GFP mutant, while abundant in interphase, displayed severely depleted levels in mitosis (**Figure 3B**). Surprisingly, unlike wild type SACK1D which binds CK1α only in mitosis, both 2xML-GFP and 1-514-GFP mutants bound CK1α regardless of cell cycle stage (**Figure 3B**). By fluorescence microscopy, a loss of spindle localized SACK1D and CK1α was observed for cells expressing either the 2xML-GFP and 1-514-GFP SACK1D mutants. In contrast, CK1α and HMMR co-localized to the mitotic spindle in cells expressing the wild type SACK1D-GFP control (**Figure 3C**). Reciprocally, guided by the Alpahfold3 prediction of the HMMR-SACK1D interaction interface (**Figure S2**), we generated a HMMR*^L37A3, F374A^*-GFP (HLF-GFP) mutant to disrupt the predicted interaction with SACK1D (**Figure 3A**). Indeed, while WT HMMR-GFP (HWT-GFP) restored in HMMR^-/-^ U2OS cells binds SACK1D, the HLF-GFP mutant abolished SACK1D binding in both asynchronous and mitotic cells (**Figure 3D**). Interestingly, we also noted that restoration of HMMR (HWT-GFP) in HMMR^-/-^ U2OS cells rescued basal SACK1D levels that were reduced in HMMR*^-/-^* cells in asynchronous conditions and fully restored the mitotic electrophoretic mobility shift of SACK1D, indicative of its mitotic hyperphosphorylation (**Figure 3D**). However, the HLF-GFP mutant was unable to rescue basal SACK1D levels, or its hyperphosphorylation in mitosis (**Figure 3D**). This is consistent with the observed rescue of SACK1D levels and mitotic phosphorylation upon restoration of HMMR-GFP and HMMRΔE4-GFP in HMMR^-/-^ cells (**Figure 2B**).

**Figure 3.**
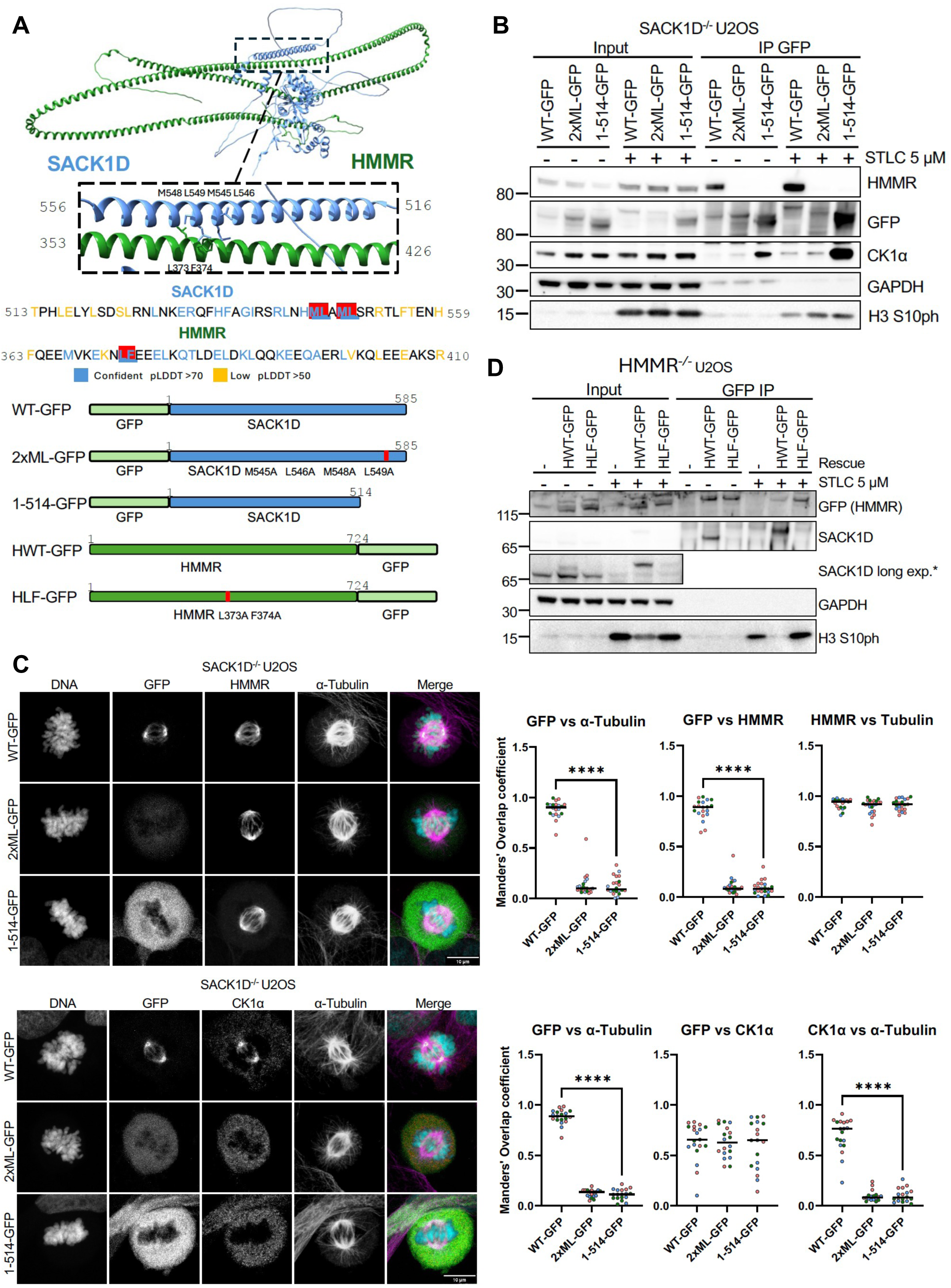
The C-terminal alpha-helix of SACK1D mediates its interaction with HMMR for spindle localization and confers mitotic specificity to its binding with CK1α. **A.** Representation of AlphaFold3 SACK1D-HMMR binding predictions with possible binding interface highlighted (top). Amino acids in the sequence coloured based on pLDDT confidence of residue’s involvement in mediating the interaction (middle). The bottom panels show schematics of the SACK1D and HMMR constructs, designed based on AlphaFold3 structural predictions. **B**. Representative western blot analysis with the indicated antibodies of lysate inputs and GFP-SACK1D IPs from SACK1D^-/-^ U2OS cells expressing the indicated SACK1D-GFP variants (2xML: SACK1D^M545A/L546A/M548A/L549A^) in asynchronous cells or those treated with STLC at 5 μM for 16 h to induce mitotic arrest. **C**. Representative confocal microscopy images of indicated GFP-SACK1D mutants expressed in SACK1D^-/-^ U2OS cells. Graphs indicate quantification of widefield immunofluorescence indicating colocalization of indicated proteins using Mander Overlap Coefficient for each protein labelled in 3 independent experiments (n>=20). Dots indicate individual cells with colours representing independent experiments (**** p<0.001). **D.** Representative western blot analysis with the indicated antibodies of lysate inputs and GFP IPs of HMMR-GFP variants (HWT: HMMR-WT; HLF: HMMR^L373A/F374A^) restored in HMMR^-/-^ U2OS cells as labelled in asynchronous cells, or those treated with STLC at 5 μM for 16 h to induce mitotic arrest.

### HMMR stabilizes full length SACK1D

To provide further evidence for the role of HMMR in recruiting SACK1D to the spindle via direct interaction with SACK1D, the HMMR binding fragment of SACK1D (aa514-585) was fused to GFP and expressed in SACK1D^-/-^ and HMMR^-/-^ U20S cells. As shown by representative confocal microscopy images, GFP-SACK1D^514-585^ restored in SACK1D^-/-^ cells is recruited to the mitotic spindle but not in HMMR^-/-^ cells (**Figure 4A**). As this provides further evidence for this region on SACK1D being responsible for HMMR interaction, to understand the extent of the role HMMR plays in SACK1D-CK1α complex formation and apparent SACK1D stabilization, WT-GFP, 2xML-GFP and 1-514-GFP were also expressed in HMMR*^-/-^* U2OS cells (**Figure 4B**). As expected, we saw no CK1α co-precipitation with WT-GFP IPs in the absence of HMMR, neither in interphase nor mitotic cells (**Figure 4B**). Surprisingly, 2xML-GFP IPs did not co-precipitate CK1α but 1-514-GFP SACK1D IPs co-precipitated CK1α robustly, independent of cell cycle stage (**Figure 4B**). On closer inspection, we noticed that expression of both WT-GFP and 2xML-GFP in HMMR*^-/-^* U2OS cells were substantially lower compared to 1-514-GFP. This reduction in abundance appeared to be further exacerbated in mitosis and suggests that HMMR stabilizes full-length SACK1D through direct interaction. Consistent with this notion, we note that HMMR*^-/-^* cells display lower levels of SACK1D protein compared to WT U2OS cells under asynchronous conditions, while in mitosis this difference in SACK1D abundance is further exacerbated as almost no SACK1D can be detected in mitotic HMMR*^-/-^* cells **(Figure 1C)**. This may also explain our previous observations of why 2xML-GFP mutants are destabilized in mitosis (**Figure 3B**) and the low abundance of WT SACK1D-GFP and 2xML-GFP in HMMR*^-/-^* cells along with their apparent further mitotic destabilization (**Figure 4B**). The evidence for HMMR regulating SACK1D stability, while compelling, is indirect and relies mostly on a single HMMR^-/-^ CRISPR clone, which may have irreversibly altered protein expression and/or signalling. To test the theory that HMMR regulates SACK1D stability directly, HMMR was depleted using siRNAs in WT U2OS cells and SACK1D abundance examined (**Figure 4C**). Indeed, the loss of HMMR leads to a robust reduction in SACK1D levels, with a further decrease in levels observed in mitotic cells (**Figure 4C**). Paradoxically, ablation of SACK1D appeared to lead to an increased accumulation of HMMR (**Figure 1D**). However, this was not reproducible when siRNAs were used to deplete SACK1D (**Figure 4D**), suggesting SACK1D is unlikely to regulate the stability of HMMR in the same way that HMMR regulates SACK1D stability. Furthermore, loss of SACK1D did not affect the rate of HMMR turnover upon mitotic exit **(EV2C)**. In WT U2OS cells, we see a robust mitotic SACK1D phosphorylation and after 3-5 hours of STLC washout, we observe almost complete and synchronous degradation of both SACK1D and HMMR (**EV2C**). In SACK1D^-/-^ cells, although there is an increase in basal and mitotic HMMR levels (a likely CRISPR artefact), the rate and extent of post-mitotic HMMR degradation did not alter substantially compared to WT cells (**EV2C**).

**Figure 4.**
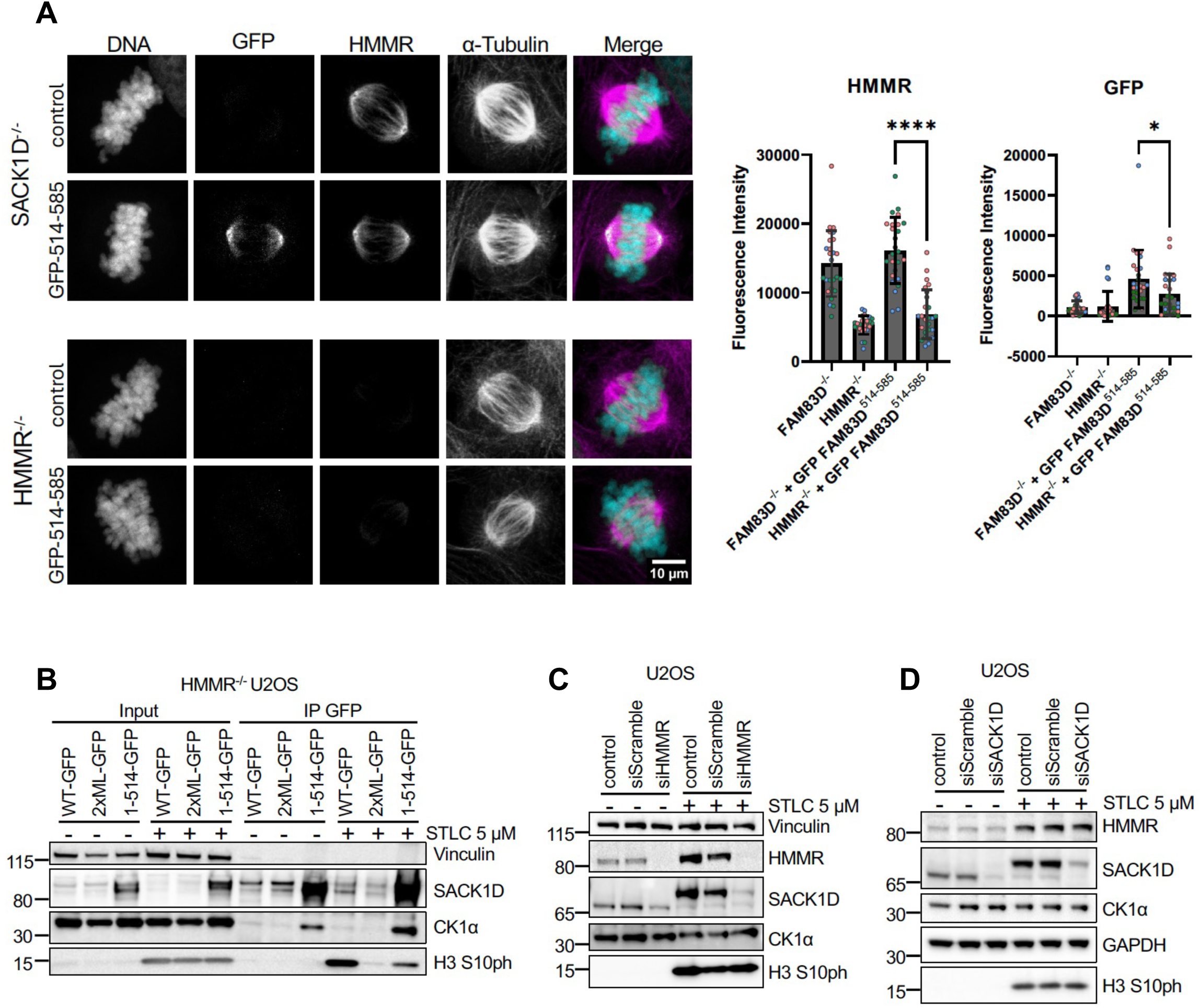
HMMR is essential for the stabilization of SACK1D, especially in mitosis. **A.** Representative confocal immunofluorescence microscopy images of SACK1D^-/-^ and HMMR^-/-^ U2OS cells expressing GFP-FAM83^514-585^. Graph indicates relative florescence intensity of indicated protein on α-tubulin representative of three independent experiments. Dots indicate individual cells with different colours representing independent experiments. (n=26, **** p<0.001) **B.** Representative western blot analysis with the indicated antibodies of lysate inputs and GFP IPs from HMMR^-/-^ U2OS cells expressing the indicated SACK1D-GFP variants in asynchronous or those treated with STLC at 5 μM for 16 h to induce mitotic arrest. **C.**Representative western blot analysis with the indicated antibodies of lysate inputs from WT U2OS cells transfected with HMMR siRNA for 24 h and lysed in asynchronous or mitotic (5 μM STLC for 16 h) conditions. **D.** Representative western blot analysis with the indicated antibodies of lysate inputs from WT U2OS cells transfected with SACK1D siRNAs for 24 h and lysed in asynchronous or mitotic (5 μM STLC for 16 h) conditions.

### Mitotic phosphorylation of SACK1D and its C-terminal α-helix signal its destruction upon mitotic exit

As shown above, and consistent with previous observations (Fulcher *et al*., 2019), SACK1D is hyperphosphorylated in mitosis. Although we found that the interaction between SACK1D and CK1α was essential to drive this mitotic SACK1D hyperphosphorylation, the precise role of CK1α in SACK1D phosphorylation and the function of these phosphorylation events remains to be determined. To more closely examine the role of CK1α and its catalytic activity on SACK1D hyperphosphorylation, we generated *CSNK1A1^-/-^* DLD-1 colorectal cells by CRISPR/Cas9 genome editing (Glennie *et al*, 2025), as repeated efforts to generate *CSNK1A1^-/-^* U2OS cells with CRISPR/Cas9 genome editing failed, potentially suggesting genetic ablation of CK1α is lethal in this background. We employed *CSNK1A1^-/-^* DLD-1 cells and stably restored the expression of either WT mCherry(mCh)-CK1α or a catalytically inactive mCh-CK1α*^N141A^*mutant (**Figure 5A)**. Unlike in WT U2OS cells where SACK1D runs at ∼75 kDa in asynchronous extracts and ∼100 kDa in mitotic cells **(Figure 1C)**, SACK1D was evident at both ∼75 kDa and ∼100 kDa in asynchronous WT DLD-1 cells, potentially suggesting a high mitotic index in asynchronous conditions **(Figure 5A)**. Despite this, in mitotic WT DLD-1 cells, only the hyperphosphorylated SACK1D migrating at ∼100 kDa was evident **(Figure 5A)**. This mobility shift of SACK1D in mitosis in WT DLD-1 cells was severely impaired in *CSNK1A1^-/-^*DLD-1 cells (**Figure 5A**). Restoration of WT mCh-CK1α in *CSNK1A1^-/-^* cells but not the catalytically inactive mCh-CK1α*^N141A^*mutant completely rescued the mitotic hyperphosphorylation of SACK1D comparable to that seen in WT DLD-1 cells (**Figure 5A**), suggesting that the catalytic activity of CK1α is essential for mitotic SACK1D hyperphosphorylation. Furthermore, we also noticed that post-mitotic destruction of SACK1D observed in WT and *CSNK1A1^-/-^* DLD-1 cells rescued with WT mCh-CK1α was impaired in *CSNK1A1^-/-^* DLD-1 cells or those rescued with the catalytically inactive mCh-CK1α*^N141A^*mutant (**Figure 5A**). These observations prompted us to hypothesize that SACK1D hyperphosphorylation by CK1α in mitosis may promote its destruction. To investigate this, we employed a phospho-proteomic analysis on mitotic SACK1D followed by site-directed mutagenesis of enriched phospho-sites (EV3A-C). This allowed us to rule out the involvement of 4 identified pS/pT residues in the observed mitotic SACK1D mobility shift (EV3A-C). Considering that the hyperphosphorylated mitotic phospho-peptide(s) responsible for the profound mitotic mobility shift may have evaded detection by mass spectrometry, potentially due to poor ionization, we focussed on a Ser-rich region of SACK1D (EV3D). We then systematically mutated 9 serine residues to alanine, either individually or collectively (EV3E). When all 9 serine residues were mutated to alanine (SACK1D-9SA), we observed a near-complete disappearance of the mitotic SACK1D hyperphosphorylation (**Figure 5C**), although none of the individual point mutations altered the mitotic mobility shift (EV3E). Interestingly, the post-mitotic destruction of SACK1D-9SA was less efficient than WT SACK1D (**Figure 5D**), suggesting that CK1α-dependent phosphorylation of SACK1D in mitosis potentially triggers its degradation upon mitotic exit. Given the reduced SACK1D abundance observed in HMMR*^-/-^* U2OS cells and the concomitant high abundance of the SACK1D^1-514^ fragment when expressed in SACK1D*^-/-^* cells, we postulated that the C-terminus of SACK1D, in addition to mediating HMMR binding, may also be crucial for its stability. Indeed, compared to WT SACK1D-GFP, which showed robust post-mitotic degradation, SACK1D^1-514^-GFP restored in SACK1D^-/-^ cells displayed an impaired post-mitotic degradation, despite its ability to bind to CK1α independently of the mitotic context (**Figure 5E**). These data suggest that the phosphorylation of the serine-rich region of the full-length SACK1D potentially triggers the recognition of the mitotic degron sequences situated at the C-terminus of SACK1D in the same region where it binds HMMR.

**Figure 5.**
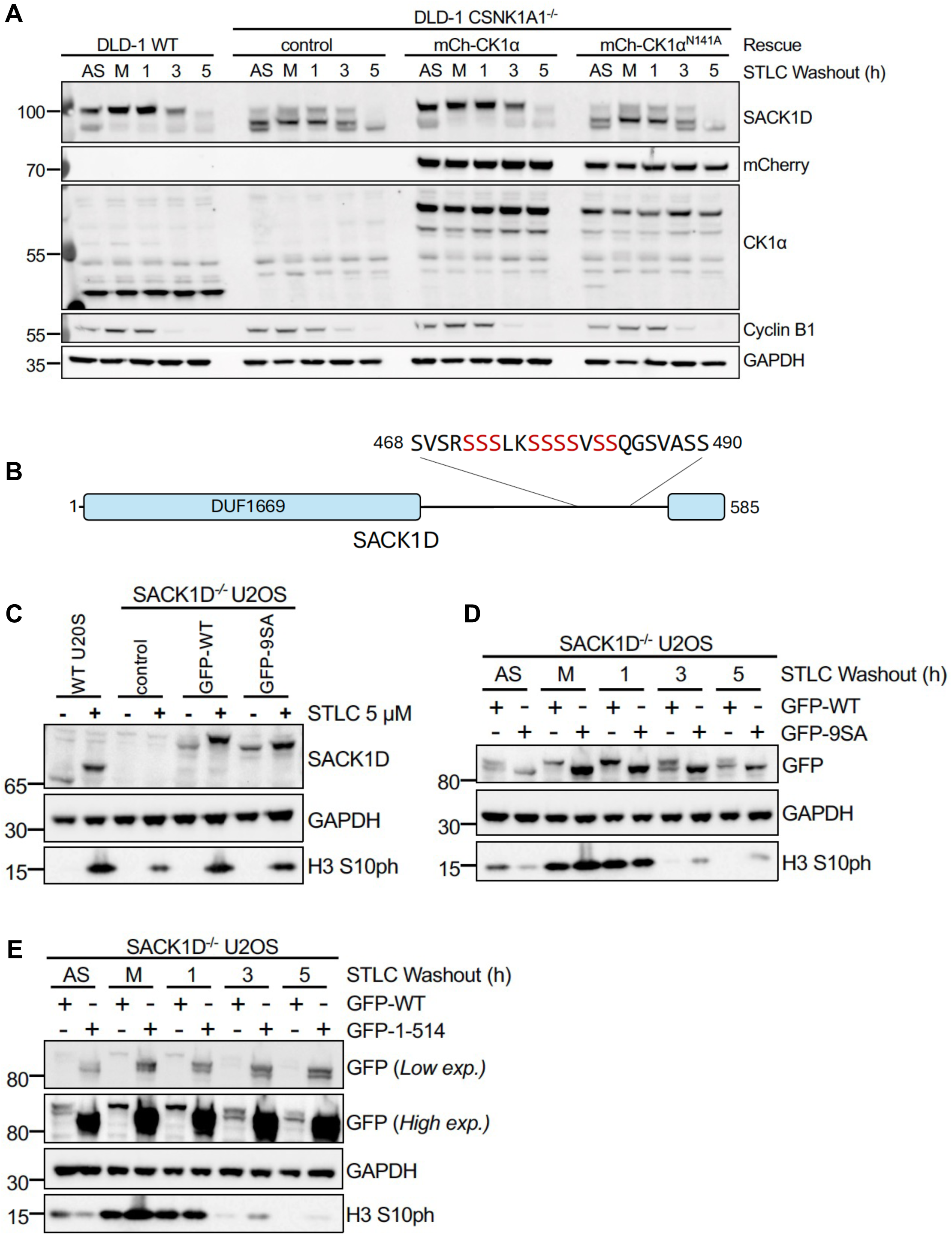
CK1α-dependant phosphorylation of SACK1D and its C-terminal α-helix mark SACK1D for rapid post-mitotic destruction. **A**. Representative western blot analysis with the indicated antibodies of lysate inputs from an STLC washout assay as indicated. WT DLD-1, *CSNK1A1^-/-^* DLD-1 or *CSNK1A1^-/-^* DLD-1 cells stably expressing either mCherry-CK1α or catalytically inactive mCherry-CK1α^N141A^ mutant were left untreated (AS) or treated with STLC at 5 μM for 16 h to induce mitotic arrest, mitotic cells isolated by shake-off and plated in fresh medium without STLC prior to lysis at the indicated time post mitotic shake-off. Cyclin B1 was employed as a control for post-mitotic degradation. **B.** Schematic of SACK1D showing the location of its serine rich region which determine the mitotic hyperphosphorylation-induced electrophoretic mobility shift by western blotting. **C.** Representative western blot analysis with the indicated antibodies of lysate inputs from WT U2OS and SACK1D^-/-^ U2OS cells expressing GFP-SACK1D or GFP-SACK1D^9SA^ (9 Ser residues indicated in B in red mutated to Ala) in asynchronous and mitotic cells (5 μM STLC for 16 h). **D.**Representative western blot analysis with the indicated antibodies of lysate inputs from an STLC washout assay (as in A) using SACK1D^-/-^ U2OS cells expressing the indicated GFP-SACK1D or GFP-SACK1D^9SA^ mutant in asynchronous and mitotic cells (5 μM STLC for 16 h). **E.**As in D except using SACK1D^-/-^ U2OS cells expressing GFP-SACK1D or GFP-SACK1D^1-514^ fragment.

## Discussion

In this study, we shed light on the intricate molecular interplay involved in the correct spatio-temporal recruitment of CK1α to the mitotic spindle in dividing cells (**Figure 6)**. We demonstrate that the microtubule-associated protein HMMR is responsible for recruiting SACK1D to microtubules and establish that a coiled-coil region comprising residues 363-410 of HMMR (specifically L373 & F374) interacts with an α-helix of SACK1D comprising residues 516-556 (specifically M545-L549). Disrupting several residues within these regions completely abolishes HMMR-SACK1D binding as well as the spindle localization of SACK1D. This is consistent with a previous study which showed that deleting a large fragment from HMMR comprising aa365-546 abolished SACK1D binding and similarly, deleting aa383-585 from SACK1D impeded interaction with HMMR (Dunsch *et al*., 2012). SACK1D recruits CK1α to the mitotic spindle (Fulcher *et al*., 2019). Ablation of HMMR from cells or restoration of SACK1D mutants that no longer bind to HMMR in SACK1D KO cells completely block the localization of CK1α to the mitotic spindle, suggesting that HMMR is crucial for the recruitment of the SACK1D-CK1α complex to the mitotic spindle. Previously, we had shown that when the spindle localization of CK1α was blocked, either via SACK1D ablation or via knocking in a SACK1D^F283A^ mutation that is incapable of binding CK1α, the cells displayed impaired spindle alignment and delayed mitotic progression (Fulcher *et al*., 2019). Consistent with this, here we find that ablation of HMMR, which leads to the absence of SACK1D-CK1α complex at the mitotic spindle, also causes misalignment of the mitotic spindles. Our data suggests that the principal role of HMMR in mitosis is to recruit the SACK1D-CK1α complex to the mitotic spindle to control proper spindle positioning.

**Figure 6.**
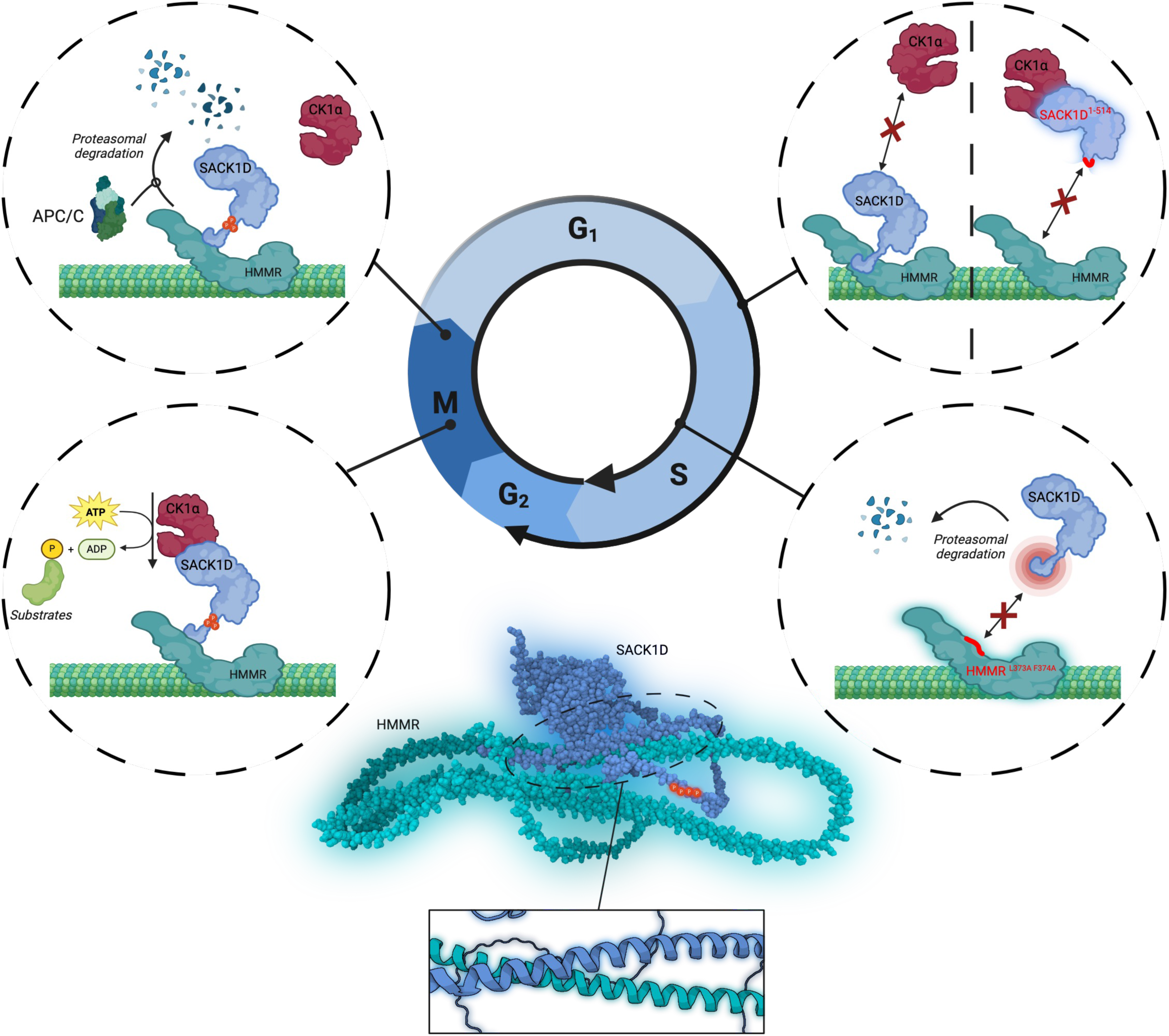
A graphical summary of key findings and the molecular insights elucidated by this study.

Peculiarly, disrupting the SACK1D-HMMR interaction by removing or mutating the C-terminal coiled-coil region of SACK1D led to its constitutive binding to CK1α, independent of the mitotic context. This further strengthens the argument that the C-terminus of SACK1D acts as its own negative regulator of CK1α binding in interphase and implies that its interaction with HMMR is integral to this negative regulatory mechanism (Fulcher *et al*., 2019). Removal of the C-terminal helix also led to stabilization of SACK1D, suggesting the presence of degron sequences at the C-terminus of SACK1D. Consistent with this notion, mutating the C-terminal helix residues (2xML->AA) from SACK1D to abolish the HMMR interaction while retaining the rest of the C-terminus, resulted in reduced stability compared to wild type SACK1D. This implies that the C-terminus of SACK1D not only contains the degron motifs but that they are protected when SACK1D is bound to HMMR. HMMR appears to be critical for the stabilization of full-length SACK1D, especially in mitosis. This means that in any context where cells lack HMMR or harbour a mutant HMMR that cannot interact with SACK1D, the spindle defects that are associated with the absence of SACK1D-CK1α complex at the spindle would also persist. However, ablation of SACK1D, which results in the absence of CK1α from the spindle, does not appear to impact HMMR stability or its post-mitotic destruction, suggesting that the SACK1D-CK1α complex does not directly regulate HMMR stability in mitosis.

We show that mitotic SACK1D hyperphosphorylation plays a role in its stability. Loss of SACK1D phosphorylation by the genetic ablation of CK1α (the kinase responsible for most of the electrophoretic mobility shift on SACK1D observed in mitosis) or mutating 9 serine residues responsible for the mitotic SACK1D mobility shift protect SACK1D from post-mitotic destruction. This is consistent with our previous observations whereby targeted dephosphorylation of SACK1D via recruitment of PPP1CA and PPP2CA, which caused a collapse of the mitotic SACK1D hyperphosphorylation electrophoretic mobility shift, impeded the rate of post-mitotic destruction of SACK1D (Simpson *et al*, 2023). Unlike SACK1D and HMMR, CK1α itself is not degraded over the course of a cell division cycle. Together, these data suggest a highly regulated mechanism by which SACK1D recruits CK1α to the mitotic spindle and ensures CK1α is not mislocalized following mitosis. Any SACK1D not bound to HMMR is degraded to potentially avoid sequestration of CK1α and any SACK1D which has been activated and binds CK1α is subsequently heavily phosphorylated and primed for degradation so that CK1α can be released back into the cytoplasmic pool and carry out other functions. In mitosis, it is through both the hyperphosphorylation of SACK1D and destruction of HMMR that SACK1D may become exposed for its destruction upon mitotic exit. Closer inspection of the C-terminal region on SACK1D reveals multiple putative D-box motifs that could potentially engage with the APC/C via Cdc20 (Davey & Morgan, 2016). Expression of the C-terminus of SACK1D containing putative D-box motifs fused to GFP leads to stable expression and spindle accumulation of GFP only in the presence of HMMR, whereas very low expression is observed in HMMR^-/-^ cells. Similarly, Alphafold3 predicts the interaction of HMMR with Cdc20 via a KEN motif providing a mechanism for destruction of HMMR by the APC/C.

CK1α is reported to control a plethora of cellular processes, including WNT, Hippo and Sonic Hedgehog (Shh) signalling, cell survival and apoptosis and, as shown here, the cell division cycle (Fulcher & Sapkota, 2020). For one protein to accomplish these diverse roles, intricate regulation of the spatio-temporal localization of CK1α is necessary and in mammalian cells, the eight members of the SACK1 domain-containing proteins play a critical role in this context (Bozatzi & Sapkota, 2018). Among the SACK1(A-H) proteins, SACK1D is unique in that its interaction with CK1α requires the mitotic context and is inhibited during interphase. This implies that the mitotic context, which requires HMMR to interact with SACK1D and deliver it to the mitotic spindle, must rapidly facilitate the SACK1 domain of SACK1D to recruit CK1α with such high affinity that both SACK1D and HMMR undergo post-mitotic degradation to release CK1α for other functions. CK1α has been shown to exist in complex with one of the other seven SACK1(A-H) proteins through their SACK1 domains (Fulcher *et al*., 2018) or GAPVD1 (Guillen *et al*, 2020; Ibrahim *et al*, 2021) under asynchronous conditions. Where the pool of CK1α that binds SACK1D at the mitotic spindle comes from remains to be determined but this is likely to be mediated by differences in affinity of the SACK1 domains of FAM83 proteins for CK1α. While this study answers many questions about the relationship between SACK1D, HMMR and CK1α, several questions remain. We show that HMMR binding is essential to the mitosis-specific recruitment of CK1α, however the exact molecular switch which allows CK1α to bind SACK1D at the G2/M transition remains elusive. The localization of CK1α to nascent spindle fibres emanating from centrosomes, combined with the reported involvement of dynein light chain 1 (DYNLL1) demonstrated by other studies, suggests that activation of the CK1α-binding interface on SACK1D may involve a complex regulatory process (Dunsch *et al*., 2012). Alternatively, post translational modification of HMMR or DYNLL1 could be sufficient to facilitate this switch. We have previously shown that the protein kinase activity of CK1α at the mitotic spindle is critical for driving proper spindle orientation and timely resolution of mitosis (Fulcher *et al*., 2019). Comprehensive assessment of the substrates modified by mitotic spindle localized CK1α remains to be elucidated. How these substrates play into the observed spindle alignment defects will further consolidate CK1α’s role as a *bona fide* mitotic kinase.

### Resource Availability

All raw mass spectrometry data acquired for this study is deposited in the PRIDE database with the accession number PXD013808 (https://www.ebi.ac.uk/pride/archive/projects/PXD013808).

### Raw Data and Code Availability

All datasets generated during this study are available at Mendeley Data link.

### Lead Contact

Further information and requests for resources and reagents should be directed to and will be fulfilled by the Lead Contact, Gopal Sapkota.

## Materials and Methods

### Cell Culture

Human osteosarcoma U2OS and colorectal adenocarcinoma DLD1 cells were grown in Dulbecco’s modified Eagle’s medium (DMEM; Gibco) containing 10% (v/v) Foetal Bovine Serum (FBS; Hyclone), penicillin (100 U/ml; Lonza), streptomycin (0.1 mg/ml; Lonza), l-glutamine (2 mM; Lonza) and Plasmocin (5ug/ml, Invivogen) and cultured at 37°C, 5% CO2 in a humidified tissue culture incubator. All cells used in this study were verified to be free from mycoplasma contamination by PCR and/or Immunofluorescence. Cells were exposed to different stimuli and compounds as described in the appropriate figure legends prior to lysis. For retroviral-based infections, cells were transduced with retroviruses as described previously (21).

### Retroviral Constructs

Recombinant DNA procedures were performed using standard protocols as described previously (Fulcher *et al*., 2018; Roth *et al*, 2020). Human wild-type SACK1D, HMMR and CSNK1A1 or appropriate mutants were sub-cloned into pBabeD vectors (pBabeD denotes a Dundee-modified version of the pBABE Puro vector). SACK1D and HMMR constructs harbour a Green Fluorescence Protein (GFP) tag at the N- and C-terminus, respectively, unless indicated otherwise. All constructs are available to request from the MRC-PPU reagents webpage (http://mrcppureagents.dundee.ac.uk), and the unique identifier (DU) numbers indicated below provide direct links to sequence information and the cloning strategy used. The following constructs were generated: pBabeD HMMR-eGFP (DU79082), pBABED HMMR^ΔK76-E91^-mCherry [Δexon4] (DU79095), pBABED HMMR (L374A F375A)-eGFP (DU79287), pBabeD GFP-SACK1D (DU28538), pBabeD GFP-SACK1D 9SA [S472A, S473A, S474A, S477A, S478A, S479A, S480A, S482A, S483A] (DU69765), pBabeD GFP SACK1D aa1-514 (DU75603), pBabeD GFP SACK1D x2ML [M545A, L546A, M548A, L549A] (DU75609), pBabeD GFP-SACK1D aa514-585 (DU75604) pBabeD mCherry-CK1α (DU69773) and pBabeD mCherry-CK1α N141A (DU69774). Constructs were sequence-verified by the DNA Sequencing Service, University of Dundee (http://www.dnaseq.co.uk). For amplification of plasmids, 1 μl of the plasmid was transformed into 30 μl of *Escherichia coli* DH5α competent bacteria (Invitrogen) on ice, incubated at 42°C for 45 s and then returned to ice for 2 min, before plating on LB-agar medium plate containing 100 μg/ml ampicillin. LB plates were inverted and incubated for 16 h at 37°C. Following incubation, a single colony was picked and used to inoculate 4 ml of LB medium containing 100 μg/ml ampicillin. Cultures were grown for 18 h at 37°C in a bacterial shaker (Infors HT). Plasmid DNA was purified using a Qiagen mini-prep kit as per the manufacturer’s instructions. The isolated DNA yield was subsequently analysed and quantified using a Nanodrop 1,000 spectrophotometer (Thermo Scientific).

### Generation of U2OS HMMR*^-/-^* Cells

To generate *HMMR*^−*/*−^ knockout by CRISPR/Cas9 genome editing, U2OS cells were transfected with vectors encoding a pair of guide RNAs (pBabeD P U6 HMMR Nter KI Sense A and pX335 HMMR Nter KI Antisense A) targeting around the first exon of *HMMR* (1 μg each) (DU60921 and DU60934). 16 h post-transfection, cells were selected in puromycin (2 μg/ml) for 48 h. The transfection process was repeated one more time. For the acquisition of single-cell clones of knockouts, single cells were isolated by fluorescence-activated cell sorting (FACS) using an Influx cell sorter (Becton Dickinson). Single cell clones were plated on individual wells of two 96-well plates, pre-coated with 1% (w/v) gelatin to help cell adherence. Viable clones were expanded, and successful knockout at the target locus was confirmed by both Western blotting and genomic DNA sequencing.

### Cell Lysis and Immunoprecipitation Assays

Media was gently aspirated, and cells lysed directly in lysis buffer (50 mM Tris–HCl pH 7.5, 150 mM NaCl, 1 mM EDTA and 1% Triton-X), supplemented with 1x cOmplete™ protease inhibitor cocktail (Roche), PhosSTOP complete phosphatase inhibitor (Roche) and Benzonase (E1014, Merck). Cell extracts were either clarified and processed immediately, or snap-frozen in liquid nitrogen and stored at −20°C. Protein concentrations were determined in a 96-well format using the BCA Gold assay (ThermoFisher).

For Immunoprecipitations (IPs), clarified extracts were normalised in lysis buffer to typically 0.75-1 mg/ml protein. After input aliquots were collected, lysates were incubated O/N at 4°C with protein G-sepharose beads coupled to the antibody of interest, on a rotating wheel. For anti-GFP IPs, GFP nanobody conjugated to Sepharose and for anti-CK1α IPs, anti-CK1α antibody-coupled sepharose beads were used, using in-house generated anti-CK1α antibody. Following incubation, beads were pelleted and were washed once in lysis buffer and 2–3 times in PBS. For elution, beads were resuspended in 1× SDS sample buffer (BioRad) and incubated at 95°C for 5 min.

### Cell Synchronization and Washout Assays

For synchronisation, cells were arrested at prometaphase with kinesin Eg5 inhibitor S-trityl-L-cysteine (STLC; 5 μM) for 16 h, before rounded/floating mitotic cells were isolated through mitotic shake-off. Collected mitotic cells were centrifuged and media aspirated before lysis in lysis buffer. Phospho-S10 Histone H3 (H3 S10ph) immunoblotting was employed as a marker of efficient mitotic arrest. For washout assays, cells that were synchronized with STLC were centrifuged and washed once in PBS before being plated in complete culture medium. Cells were then collected at indicated time points by media centrifugation at early time points where cells were still in suspension or direct lysis on plates where cells had attached.

### SDS-PAGE and Western Blot

Reduced protein extracts (typically 10–20 μg protein, Lamelli with DTT) or IPs were resolved on 4–12% NuPAGE bis-tris precast gradient gels (Invitrogen) by electrophoresis. Separated proteins were subsequently transferred onto nitrocellulose membranes (Amersham), before membranes were blocked in 5% (w/v) non-fat milk powder (Marvel) in TBS-T (50 mM Tris–HCl pH 7.5, 150 mM NaCl, 0.2% (v/v) Tween-20) and incubated overnight at 4°C in 5% milk TBS-T with the appropriate primary antibody. Membranes were then washed for 3 × 10 min with TBS-T before incubating with HRP-conjugated secondary antibodies in 5% milk TBS-T for 1 h at RT. Membranes were then washed for 3 × 10 min with TBS-T before detection with enhanced chemiluminescence reagent (Invitrogen) and imaged using the ChemiDoc™ system (Bio-Rad).

### siRNA Transfection

U2OS cells were transfected with esiRNAs (600ng in 2ml in a well of a 6 well plate) targeting HMMR (EHU060291, Merck), FAM83D/SACK1D (EHU084291, Merck) or the universal negative control 1 (SIC001, Merck) using Lipofectamine RNAiMax Reagent (13778075, Thermo Fisher). Lipid–siRNA mixtures were incubated in Opti-MEM (Gibco) for 5 min before being added dropwise to cells. Cells were collected for analysis after 24 h with co-treatment with STLC (5 μM) or DMSO added for 16 h before lysis as required.

### Confocal and Widefield Immunofluorescence Microscopy

Cells were seeded onto glass coverslips and treated/transfected as described above or in figure legends. Media was gently aspirated before fixing in 4% (w/v) paraformaldehyde (PFA) in x1 PHEM buffer (60 mM PIPES, 25 mM HEPES, 10 mM EGTA, and 4 mM MgSO4·7H20) for 10 min at 37°C. Cells were washed twice in PBS, followed by incubation in PBS at 4°C or stained immediately. Cells were permeabilised for 5 min in 0.1% Triton in PBS. Following permeabilization, cells were blocked in PBS containing 5% (w/v) BSA for 1 h. Where appropriate, coverslips were then incubated with primary antibody in PBS/BSA 0.1% Triton (at dilutions indicated in the antibody table) at 37°C for 1 h. Cells were washed for a minimum of 3 × 10 min in PBS/BSA before incubation with the secondary AlexaFluor-conjugated antibody in PBS/BSA 0.1% Triton (1:1000) for 60 min at RT protected from light. Coverslips were subsequently washed for 1 × 10 min each in PBS 0.1% Triton, PBS and MilliQ Water, then dried and mounted on glass microscopy slides using ProLong® Glass anti-fade reagent with NucBlue (Life Technologies). Coverslips were left to set overnight at 4°C before analysis on a Leica Stellaris 8 Falcon FLIM confocal microscope using an HC PL APO 63x/1.40 OIL CS2 lens. Alternatively, slides were imaged on a DeltaVison Elite widefield microscope using x100 PlanApoN 1.42 NA objective at 1 × 1 binning and a CoolSNAP HQ camera (Photometrics).

### Immunofluorescence Microscopy Analysis

All image analysis was performed in FIJI. For quantification of CK1α, SACK1D and HMMR on the mitotic spindle for siRNA studies (Figure 1D), the mean pixel intensities of proteins of interest were calculated by designating a circular region of interest (ROI) around the DNA of the cell and subtracting the mean pixel intensity of the background (ROI) outside the first region but inside the cell. For quantification of HMMR and CK1α in HMMR rescue experiments (Figure 2A) and GFP-SACK1D^514-585^ rescue experiments (Figure 4A) mean pixel in intensity was obtained for an ROI determined by α-tubulin staining (MaxEntropy Threshold) and subtracting the mean pixel intensity of the background (ROI) outside the first region but inside the cell. For colocalization assays (Figure 3C), Manders overlap coefficients were obtained with the JACoP plugin for proteins as indicated. Macros used are supplied in Appendix 1.

### Mitotic Spindle Alignment Assay

To determine mitotic spindle orientation, cells were stained with pericentrin antibody to indicate centrosomes and the position of each centrosome relative to the other recorded in prometaphase-metaphase cells. A minimum of 24 cells per condition were recorded spread over three independent experiments.

### Alphafold3 Analysis

The canonical sequence of human HMMR and FAM83D/SACK1D obtained from UNIPROT were uploaded to the Alphafold3 webserver at https://alphafoldserver.com/. The highest confidence prediction was then cross referenced with other predictions provided in Chimera X (UCSC) and predicted interactions determined by PAE analysis of bonds forming between predicted intermolecular contacts.

### Statistical Analysis and Reproducibility

Statistical analysis was carried out in GraphPad (Prism) and was used to generate plots and analyse data by unpaired Student’s t-test or one-way ANOVA as appropriate following normality tests. A P-value of < 0.05 was deemed significant. Unless indicated otherwise, all experiments are representative of at least three independent biological replicates. Where appropriate, the data from all three replicates is deposited in at Mendeley Data linked to this manuscript.

### Antibodies

The antibodies used in this study are listed in Table 1. The source, catalogue number and dilution factor used for each antibody are provided. The in-house antibodies were generated by the MRC PPU Reagents and Services, University of Dundee as described previously (22). For HRP-coupled secondary antibodies, goat anti-rabbit-IgG (cat.: 7074, 1:1000) was from CST, and rabbit anti-sheep-IgG (cat.: 31480, 1:1,000) and goat anti-mouse-IgG (cat.: 31430, 1:1,000) were from Thermo Fisher Scientific.

**Table 1.** Primary Antibodies used in this study.

| Antigen | Source | Cat. Number | Dilution | Figure | Species |
| --- | --- | --- | --- | --- | --- |
| GFP | MBL | 589 | 1:1000<br>(WB) | 4B | Rabbit |
| GFP | DSTT<br>(University of Dundee) | S268B (Bleed 4) | 1:1000<br>(WB)<br>1:500 (IF) | 1A, 3B, 5D&E | Sheep |
| mCherry | Abcam | ab213511 | 1:1000<br>(WB)<br>1:500 (IF) | 1A, 2A & 5A | Rabbit |
| α-Tubulin | Invitrogen | MA1-80189 | 1:1000<br>(WB) | 1A&B, 2B&D & 3C | Rat |
| Vinculin | CST | 13901S | 1:5000<br>(WB) | 1C, 2C, 3E, 4C&E | Rabbit |
| HMMR | Santa Cruz | Sc-515221 | 1:1000(WB)<br>1:500 (IF) | 1B&C, 2A, B&C,<br>3B&C, 4B, C, D & E, | Mouse |
| SACK1D | DSTT<br>(University of Dundee) | SA102 | 1:1000<br>(WB)<br>1:500 (IF) | 1C, 2C, 3E, 4B, C,<br>D&E, 5A, & C & S3B | Sheep |
| Ck1α | DSTT<br>(University of Dundee) | SA527 Bleed 3 | 1:1000<br>(WB)<br>1ug/ml (IP) | 1C, 2C, 3B&E,<br>4C&D & 5A | Sheep |
| Ck1α | DSTT<br>(University of Dundee) | SA527 Bleed 8 | 1:500 (IF) | 2B, & 3C | Sheep |
| H3 S10ph | CST | 9701S | 1:1000<br>(WB) | 1C, 2C, 3E, 4B, C,<br>D&E & 5C, D&E | Rabbit |
| Pericentrin | Invitrogen | PA5-54109 | 1:50 (IF) | 1B & 2D | Rabbit |
| Cyclin B1 | CST | 12231S | 1:1000<br>(WB) | 5A | Rabbit |
| GAPDH<br>(HRP<br>conjugated) | ProteinTech | HRP-60004 | 1:5000<br>(WB) | 3B&D, 4D,<br>5A,C,D&E, & EV3 | Rabbit |

**Table 2.** Secondary Antibodies used for immunofluorescence in this study.

| Target Species | Raised Species | Wavelength | Source | Catalog Number | Dilution Factor |
| --- | --- | --- | --- | --- | --- |
| Rabbit | Goat | 594nM | Invitrogen | A11012 | 1:1000 |
| Rat | Donkey | 647nM | Invitrogen | A78947 | 1:1000 |
| Sheep | Donkey | 488nM | Invitrogen | A11015 | 1:1000 |
| Sheep | Donkey | 594nM | Invitrogen | A11016 | 1:1000 |
| Mouse | Donkey | 488nM | Invitrogen | A21202 | 1:1000 |
| Mouse | Goat | 594nM | Invitrogen | A11005 | 1:1000 |
| Mouse | Goat | 647nM | Invitrogen | A32728 | 1:1000 |

## Author Contributions

TNC performed most of the experiments, collected & analysed data and contributed to the writing of the manuscript. NKN, LJF, KD & SB performed some experiments and collected & analysed data. NTW performed cloning. TJM designed the strategies for and generated the CRISPR/Cas9 constructs used in this study. GPS conceived the project, analysed data, and contributed to the writing of the manuscript.

## Acknowledgements

GPS is supported by the UKRI Medical Research Council (grant MC_UU_00018/6 and MC_UU_00038/6) and Boehringer Ingelheim through the Division of Signal Transduction Therapy. We thank the Sapkota lab members for critical appraisal of the data. We thank E. Allen, A. Muir, S. Dalglish, E. Webster and J. Stark for help and assistance with tissue culture, the staff at the DNA Sequencing services (School of Life Sciences, University of Dundee), and the cloning and antibody teams within the MRC-PPU Reagents and Services (University of Dundee, coordinated by J. Hastie). We thank the staff at the flow cytometry facility (School of Life Sciences, University of Dundee) for their invaluable support. We thank the MRC PPU mass spectrometry service team for their help with the project. We thank members of the Dundee Imaging Facility (University of Dundee, coordinated by P. Appleton) for microscopy support.

## Expanded View Figure Legends

**Expanded View 1.**
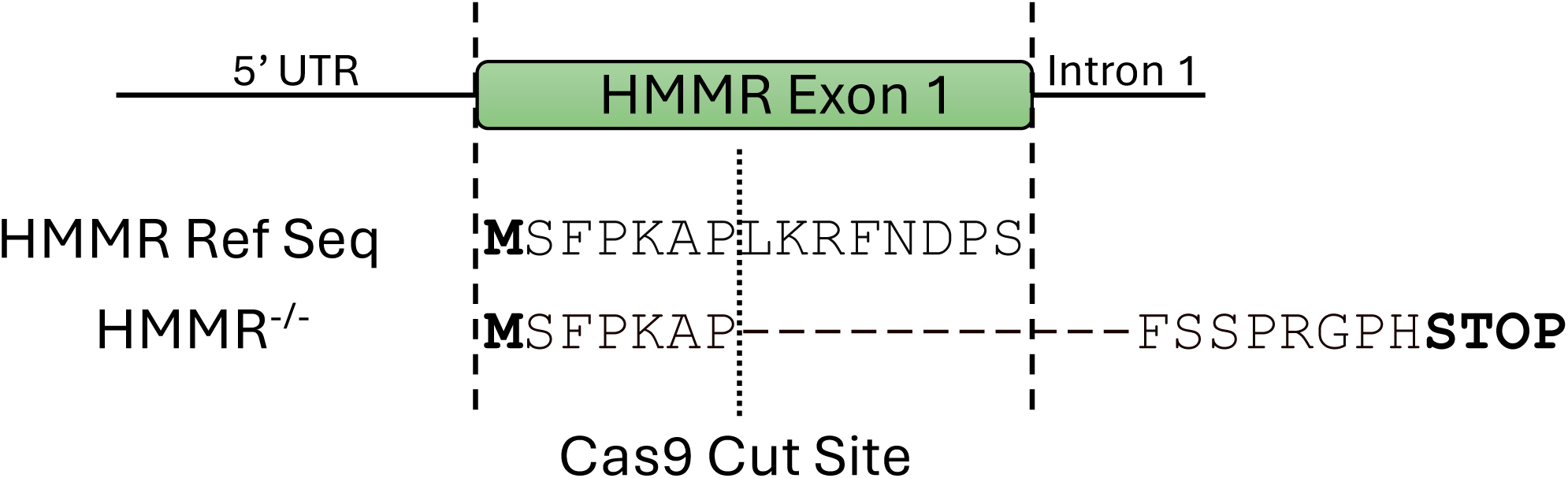
Schematic of the sequencing results confirming a deletion in exon 1 of HMMR.

**Expanded View 2.**
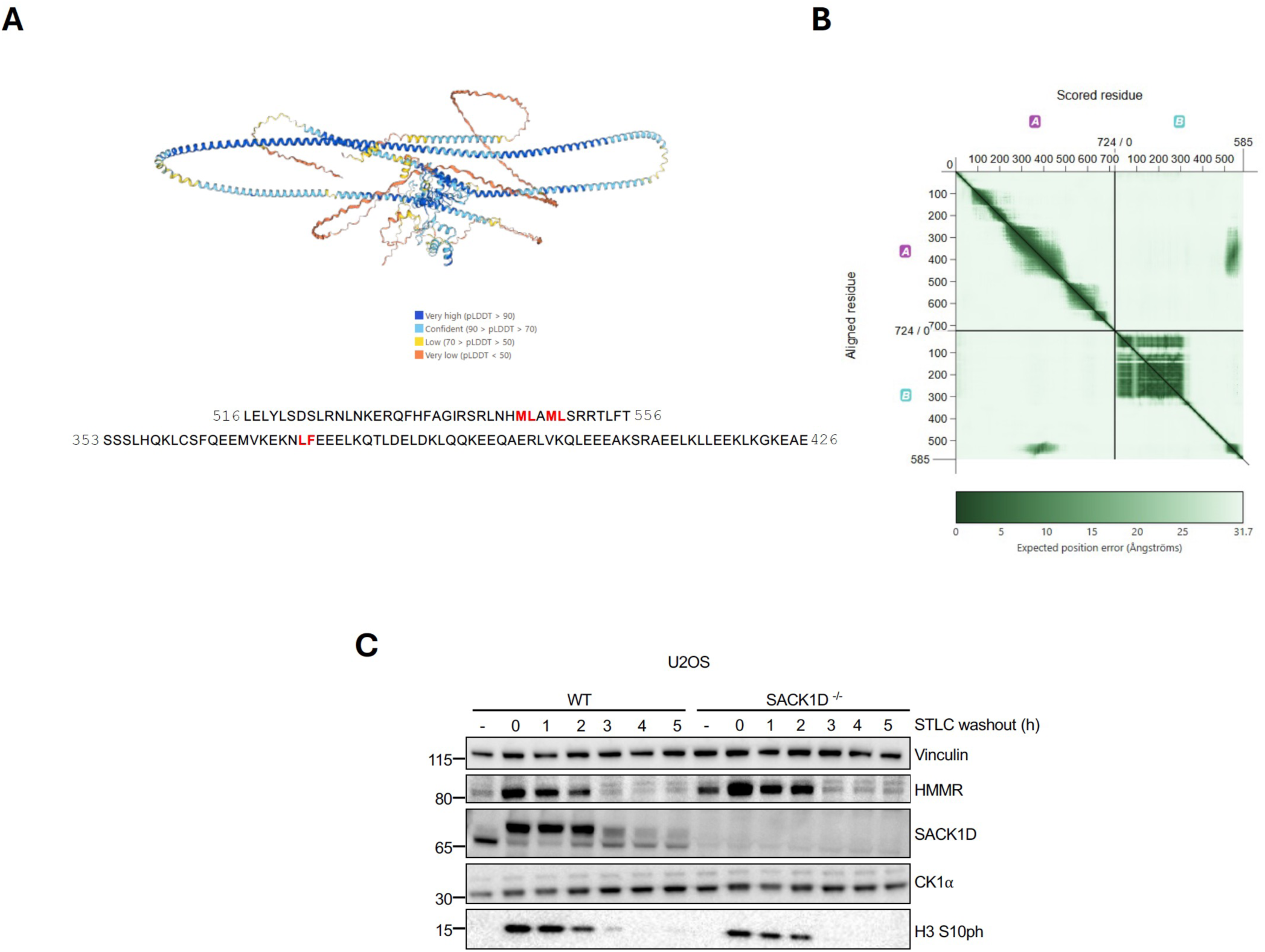
Alphafold3 prediction of SACK1D-HMMR complex depicting the binding interface. **A.** 3D representation of the predicted structure of SACK1D bound to HMMR. Colours indicate predicted Local Distance Difference Test values according to the legend provided. Amino acid sequences from SACK1D (top) and HMMR (bottom) at the interface, with the residues predicted to make molecular contacts highlighted in red. **B.**Predicted aligned error (PAE) plot of SACK1D against HMMR. **C.**Representative western blot analysis with the indicated antibodies of lysate inputs from an STLC washout assay performed in WT U2OS and SACK1D^-/-^ U2OS cells and lysed at indicated timepoints (h) following mitotic arrest.

**Expanded View 3.**
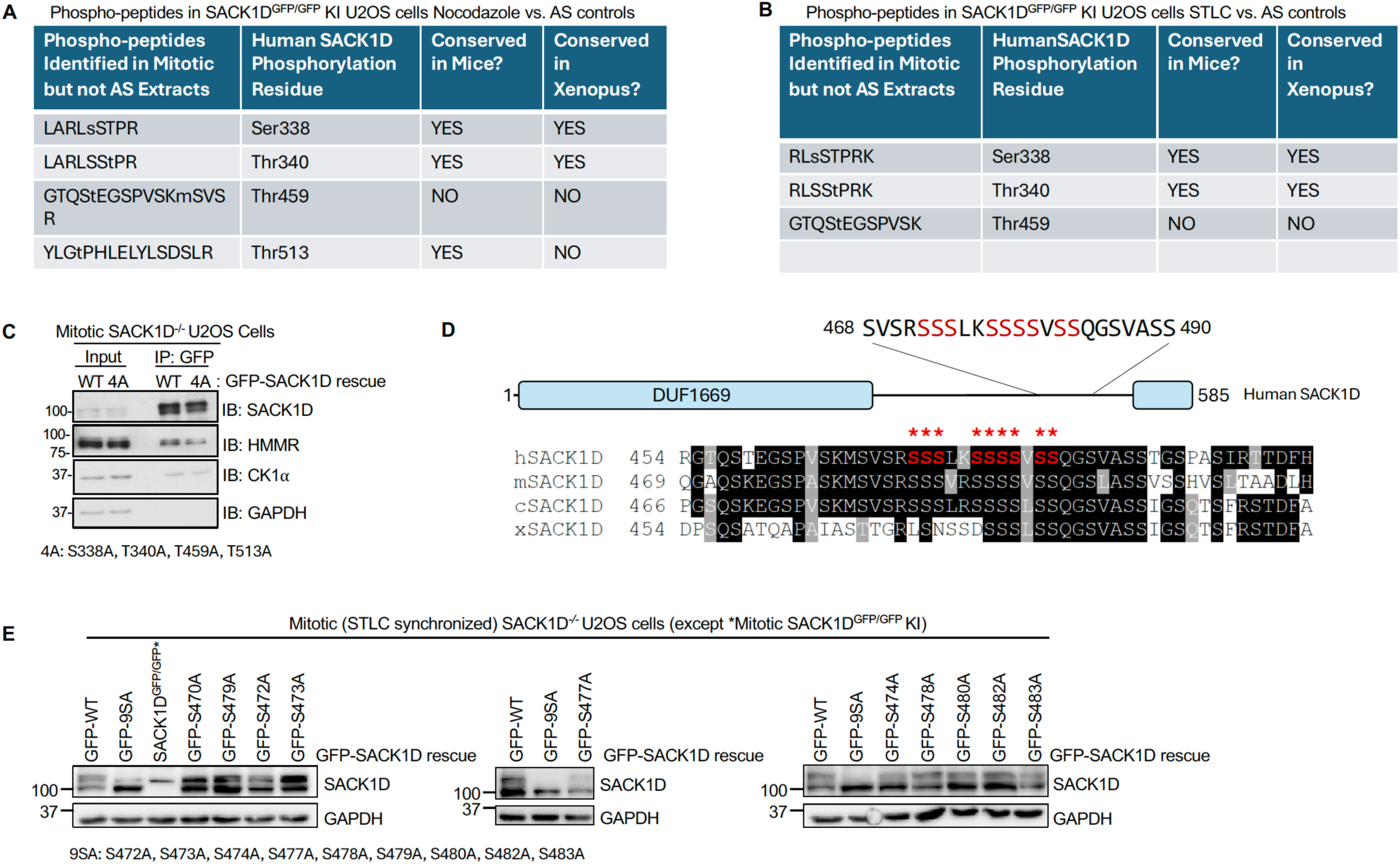
Identification of the phospho-sites on SACK1D responsible for the mitotic electrophoretic mobility shift. **A.** Table showing the phospho-peptides on SACK1D identified by mass-spectrometry in anti-GFP IPs from nocodazole-synchronized mitotic SACK1D^GFP/GFP^ knockin U2OS cell extracts that were absent in asynchronous (AS) extracts. The residue numbers corresponding to human SACK1D and whether they are conserved in mouse and *Xenopus* are indicated. **B.** As in (A), except the cells were synchronised in mitosis by treating them with STLC. **C.** The 4 phospho-residues identified in (A) were mutated to Ala (4A) and restored in SACK1D^-/-^ U2OS cells as indicated. STLC-synchronized mitotic extracts or anti-GFP IPs were subjected to western blot analysis with the indicated antibodies. The GFP-SACK1D-4A mutant did not alter the mitotic electrophoretic mobility shift of SACK1D compared to GFP-SACK1D-WT suggesting there are more mitotic phospho-sites on SACK1D that were not identified by mass-spectrometry. **D.** Schematic showing the location of the conserved serine-rich cluster region on SACK1D that we considered as a potential region that could be hyperphosphorylated in mitosis to account for the electrophoretic mobility shift. The indicated 9 Ser residues were chosen to mutate to Ala either individually or collectively. Sequence alignment of this region with human, mouse, chicken and Xenopus SACK1D was performed by Clustal Omega and image generated using BoxShade Server. **E.** Western blot analysis of single Ser to Ala point mutations and all 9 Ser to Ala (9SA) GFP-SACK1D mutants showing that only 9SA mutations result in the complete collapse of the mitotic SACK1D band shift. All mutants were retrovirally transduced in SACK1D^-/-^ U2OS cells. * Indicates U2OS SACK1D^GFP/GFP^ knockin mitotic extract included as a positive control for the observed electrophoretic mobility shift.

